# Taming Darwin’s conundrum: On the role of phylogenetic and functional differentiation in plant naturalization on oceanic islands

**DOI:** 10.1101/2025.04.09.648018

**Authors:** Yurena Arjona, Louis S. Jay-García, J. Alfredo Reyes-Betancort, Javier Morente-López, Marcos Salas-Pascual, Agustín Naranjo-Cigala, Guillermo Sicilia-Pasos, Miguel A. Padrón-Mederos, Raúl Orihuela-Rivero, Rafaela González-Montelongo, Jairo Patiño

## Abstract

- Biological invasions are a major threat to global biodiversity, particularly on oceanic islands and their unique and fragile biotas. However, the complexity of the processes underlying invasions has prevented the identification of general patterns explaining why some plant species successfully establish and spread beyond their native ranges.
- We investigated Darwin’s naturalization conundrum in the most species-rich angiosperm family of the Canary Islands, considering two invasion stages across two spatial scales. Using a high-resolution phylogeny covering the Asteraceae species from the archipelago and functional traits measured from fresh specimens, we calculated phylogenetic and functional distances of invasive and noninvasive introduced species to natives.
- Our findings reveal that successful introduced Asteraceae species were phylogenetically distant from native relatives, with invasive species even more distantly related than noninvasive introduced species to their closest native relative. Functionally, noninvasive species were distinct, while invasive species showed unexpected similarity to native species. These patterns were consistent across spatial scales.
- This study underscores the importance of a multidimensional approach to understanding plant invasion success. Identifying key patterns driving biological invasions is crucial for developing effective strategies against future ecological disruptions on islands.

## Introduction

The human-mediated introduction of species beyond their native ranges, followed by their successful establishment in the wild, is reshaping global biodiversity and biogeography at an unprecedented pace (van Kleunen *et al*., 2015; Turbelin *et al*., 2017; IPBES, 2019; Pyšek *et al*., 2020). This is expected to continue and even accelerate in the future, as the number of alien species is projected to increase due to multiple anthropogenic drivers (Seebens *et al*., 2017, 2018, 2021), including climate change, land use modifications, and the expansion and technification of human mobility and trade (Banks *et al*., 2015; Hulme, 2021; Liu *et al*., 2024; Bradley *et al*., 2024). These ongoing trends are expected to exacerbate ecological and socioeconomic impacts worldwide, with one of the most severe consequences being the extinction of native species (Bellard *et al*., 2016; Pyšek *et al*., 2020; Orihuela-Rivero *et al*., 2025).

Island ecosystems are particularly vulnerable to biological invasions (Russell & Kueffer, 2019; Fernández-Palacios *et al*., 2021). Of the global biodiversity loss driven by invasive species, 86% of the recorded extinctions were island endemics (Bellard *et al*., 2016). This heightened vulnerability is largely due to the unique geological and ecological characteristics of islands, which support exceptionally high levels of endemic species despite their small land area (Kier *et al*., 2009; Fernández-Palacios *et al*., 2021; Schrader *et al*., 2024). While islands are renowned for their rich native and endemic biodiversity, they are also among the most invaded territories on Earth, serving as hotspots for alien species (Moser *et al*., 2018; Essl *et al*., 2019; Russell & Kaiser-Bunbury, 2019). The outstanding contribution of islands to global biodiversity and their highly threatened situation justify the urgent need to deepen our understanding of invasion processes driving island biodiversity loss.

A wide range of theories and hypotheses have been proposed to explain plant invasions. However, the ecological and evolutionary drivers of biological invasions are far from uniform.These forces operate along complex gradients and vary depending on environmental context, making generalizations difficult (Theoharides & Dukes, 2007; Dai *et al*., 2020; Daly *et al*., 2023; Gioria *et al*., 2023). One of the most significant gradients concerns the invasion process itself, which has been defined as a three-stage continuum comprising introduction, naturalization and invasion (Richardson *et al*., 2000; Blackburn *et al*., 2011). Following human-mediated transport, species are introduced in new regions outside their native range. If conditions allow, some of these species establish self-sustaining populations, this is, they become naturalized in the recipient areas. Subsequently, some species may spread into (semi)natural ecosystems, potentially causing biodiversity loss and ecosystem disruption, thus reaching the final invasion stage of the continuum. Identifying the factors that govern each stage remains a challenge, as the spatio-temporal invasion dynamics can differ across spatial scales, taxa and geographic regions (Pyšek & Hulme, 2005; Theoharides & Dukes, 2007).

Inherent characteristics of the species may favour their progression through the invasion continuum (Daly *et al*., 2023; Gioria *et al*., 2023). Functional traits related to plant reproductive strategy, growth and longevity have been reported to play a crucial role in determining whether a species successfully naturalizes and spreads (Pyšek & Richardson, 2007). However, patterns remain inconsistent across studies, highlighting the strong context dependency of these trait-based relationships (van Kleunen *et al*., 2010; Richardson & Pyšek, 2012; Catford *et al*., 2019; Daly *et al*., 2023). The characteristics of the recipient native community have also been found determinant of the success of introduced species, as they relate to its vulnerability to invasion (Catford *et al*., 2019; Daly *et al*., 2023; Gioria *et al*., 2023). Ultimately, the interaction between the introduced species and the recipient community determines invasion success, making it crucial to assess the divergence between native and introduced species (Carboni *et al*., 2016; Catford *et al*., 2019; Renault *et al*., 2022). Functional traits provide insight into a species’ ecological strategy, reflecting both abiotic and biotic interactions with other species and the environment (Violle *et al*., 2007; Cadotte *et al*., 2011). In addition, evolutionary history and relatedness can constrain trait variation and shape species’ environmental affinities (Prinzing *et al*., 2001; Davies *et al*., 2013). Integrating phylogenetic and functional relationships between introduced species and recipient native communities will improve our understanding of the mechanisms driving biological invasions.

Building on this framework, the role of the relationship between introduced species and the recipient community in invasion success has been debated since Charles Darwin first questioned why some introduced species successfully naturalize in new environments while others do not (Darwin, 1859). He discussed two seemingly opposing hypotheses: the naturalization and the pre-adaptation hypotheses. Darwin’s naturalization hypothesis posits that species distantly related to, and thus functionally different from, the native flora are more likely to naturalize. This hypothesis is based on the idea that successful introduced species occupy different ecological niches and avoid direct competition with native species (biotic filtering). In contrast, Darwin’s pre-adaptation hypothesis suggests that introduced species closely related to native taxa are more likely to become established because their shared evolutionary history provides adaptations suited to local environmental conditions. In this second hypothesis, the environmental filtering is therefore dominant in driving invasion success. The apparent contradiction between these perspectives, often referred to as Darwin’s naturalization conundrum (Diez *et al*., 2008), has fueled extensive research, with studies supporting one, the other, or both hypotheses (Thuiller *et al*., 2010; Cadotte *et al*., 2018).

The ongoing debate surrounding Darwin’s hypotheses largely stems from the heterogeneity of study designs and the scale at which invasion processes are examined. At small spatial scales, where interspecific competition is more intense, the naturalization hypothesis is more likely to hold, as newly established species are expected to minimize the competition by differing from the recipient community. Conversely, at larger spatial scales, where species do not strictly coexist, the pre-adaptation hypothesis may be more applicable, as shared key adaptations between invaders and the recipient community will allow them to overcome the dominant environmental filter (Thuiller *et al*., 2010; Cadotte *et al*., 2018; Bach *et al*., 2022). The temporal dimension of the invasion process further complicates testing Darwin’s naturalization conundrum. Over time, introduced species may intensify competition with native taxa, sometimes leading to the local decline or extinction of their closest relatives.

However, long-term studies tracking the evolution of invasion remain scarce (but see Li *et al*., 2015; Catford *et al*., 2019). In turn, most studies compare naturalized and invasive species, treating them as distinct stages of the invasion continuum (Strauss *et al*., 2006; Schaefer *et al*., 2011; Park & Potter, 2013; Omer *et al*., 2022). The relative support for Darwin’s hypotheses may shift depending on the invasion stage under consideration (Li *et al*., 2015; Cadotte *et al*., 2018; Omer *et al*., 2022). Further complicating these analyses are methodological challenges, such as the phylogenetic resolution or the assumption that closely related species share similar ecological traits, both of which can confound interpretations of Darwin’s hypotheses (Park & Potter, 2013; Marx *et al*., 2016; Cadotte *et al*., 2018). Addressing adequately these issues is essential for refining our understanding of the mechanisms driving invasion success.

In the present study, we investigate Darwin’s naturalization conundrum by identifying key patterns driving the success of introduced Asteraceae species in the Canary Islands. Asteraceae is among the most diverse plant families across oceanic archipelagos worldwide (Roeble *et al*., 2024), making it an ideal system for testing hypotheses related to invasion success. We examine functional and phylogenetic divergence between introduced and native species at two spatial scales, archipelago-wide *versus* island-specific, comparing naturalized and invasive species as distinct stages of the invasion continuum (Richardson *et al*., 2000). We construct a high-resolution phylogenetic tree using over 1,000 nuclear genes per species, allowing for precise measurements of evolutionary distances between species. We also calculate a field-based multi-trait functional distance, avoiding the well-documented issue of assuming a strict correlation between phylogenetic and ecological similarity (Pavoine & Bonsall, 2011).

Specifically, our goals are to: (1) assess the phylogenetic and functional distances between naturalized (noninvasive) and native species, and compare them with those of invasive species; (2) determine how these distances deviate from random expectations; and (3) evaluate whether patterns differ between the archipelago and individual islands. Given that most introduced species in the Canary Islands originate from the Neotropics (Morente-López *et al*., 2023), whereas the native flora is largely of Mediterranean origin (Carine *et al*., 2010; but see Fernández-Palacios *et al*., 2024), we expect both noninvasive and invasive introduced species to be phylogenetically distant from native taxa. Furthermore, we anticipate a degree of decoupling between phylogenetic and functional distances, largely due to evolutionary radiations within the family. For instance, species from recent radiations may be closely related yet functionally divergent if they have adapted to distinct ecological niches (Bach *et al*., 2022). Finally, following Bach *et al*. (2022), we hypothesize that the pre-adaptation hypothesis will hold at broader scales, across the entire archipelago and larger islands, where environmental filtering plays a dominant role. In contrast, Darwin’s naturalization hypothesis is likely to be more relevant on smaller islands, where competition is stronger. This study represents the first high-resolution phylogenetic and functional analysis of Darwin’s naturalization conundrum in the flora of the Canary Islands, providing novel insights into the factors shaping plant invasions on islands.

## Material and Methods

### Study system

The Canary Islands are an oceanic archipelago, located in the Atlantic Ocean about 96 km off the southern coast of Morocco. The archipelago consists of seven large islands (> 100 km^2^), one smaller island (> 20 km^2^) and several islets, all of volcanic origin. It harbors 2,320 species of vascular flora with a high percentage of endemic species (25.2%), but an even higher percentage of introduced species (39%) (Biodiversity Data Bank of the Canary Islands, https://www.biodiversidadcanarias.es/biota/; accessed on the 10^th^ of December 2024).

Asteraceae is the most diverse family of flowering plants, comprising 314 species and a notably high amount of endemics (44%), including highly radiated genera such as *Sonchus*, *Argyranthemum* and *Cheirolophus* (Santiago & Kim, 2009; Vitales *et al*., 2014; White *et al*., 2020). Approximately 28% of the Asteraceae species present in the Canary Islands have been introduced by human means. Some of these species are causing severe ecological impacts. For example, *Ageratina adenophora* (Spreng.) R. M. King & H. Rob. and *A. riparia* (Regel.) R. M. King & H. Rob. are highly invasive weeds in several countries across Asia, Oceania, Africa and Europe (Poudel *et al*., 2019). Other invasive species, despite having been recorded in the archipelago only recently, are rapidly spreading and exerting significant ecological pressure. Examples include *Pluchea ovalis* (Pers.) DC. (Padrón-Mederos *et al*., 2009; Verloove & Reyes-Betancort, 2011; García-Alvarado *et al*., 2025) and *Youngia japonica* (L.) DC. (Siverio Núñez *et al*., 2013; Verloove, 2017). Given the Canary Islands’ unique and fragile biodiversity, along with the high costs associated with managing invasive species, it is crucial to understand the invasion process and the factors driving it. Such knowledge is essential for developing effective strategies to prevent future ecological disruptions (Otto *et al*., 2014; Steinbauer *et al*., 2017; Fernández-Palacios *et al*., 2023).

### Sampling

We compiled a checklist of the Asteraceae species present in the Canary Islands extracted from the Biodiversity Data Bank of the Canary Islands (BIOTA, https://www.biodiversidadcanarias.es/biota/; accessed for the first time in October 2020 with posterior refinements, see Supporting Information Table S1). Based on this checklist and recorded species distribution, we conducted fieldwork on all the islands of the archipelago to collect at least one fresh sample per species (Fig. 1). The number of fresh leaves collected per species depended on leaf size, with leaves sampled separately for DNA extraction and trait measurements. Whenever possible, a voucher specimen was collected and deposited in the herbarium of the Jardín de Aclimatación de La Orotava (ORT) or the institutional herbarium of the University of La Laguna (TFC) (see Supporting Information Table S2). For DNA extraction, fresh leaves were dried in silica gel; while for trait measurements, fresh leaves were introduced in plastic zip bags and carried to the laboratory for immediate processing.

**Figure 1.**
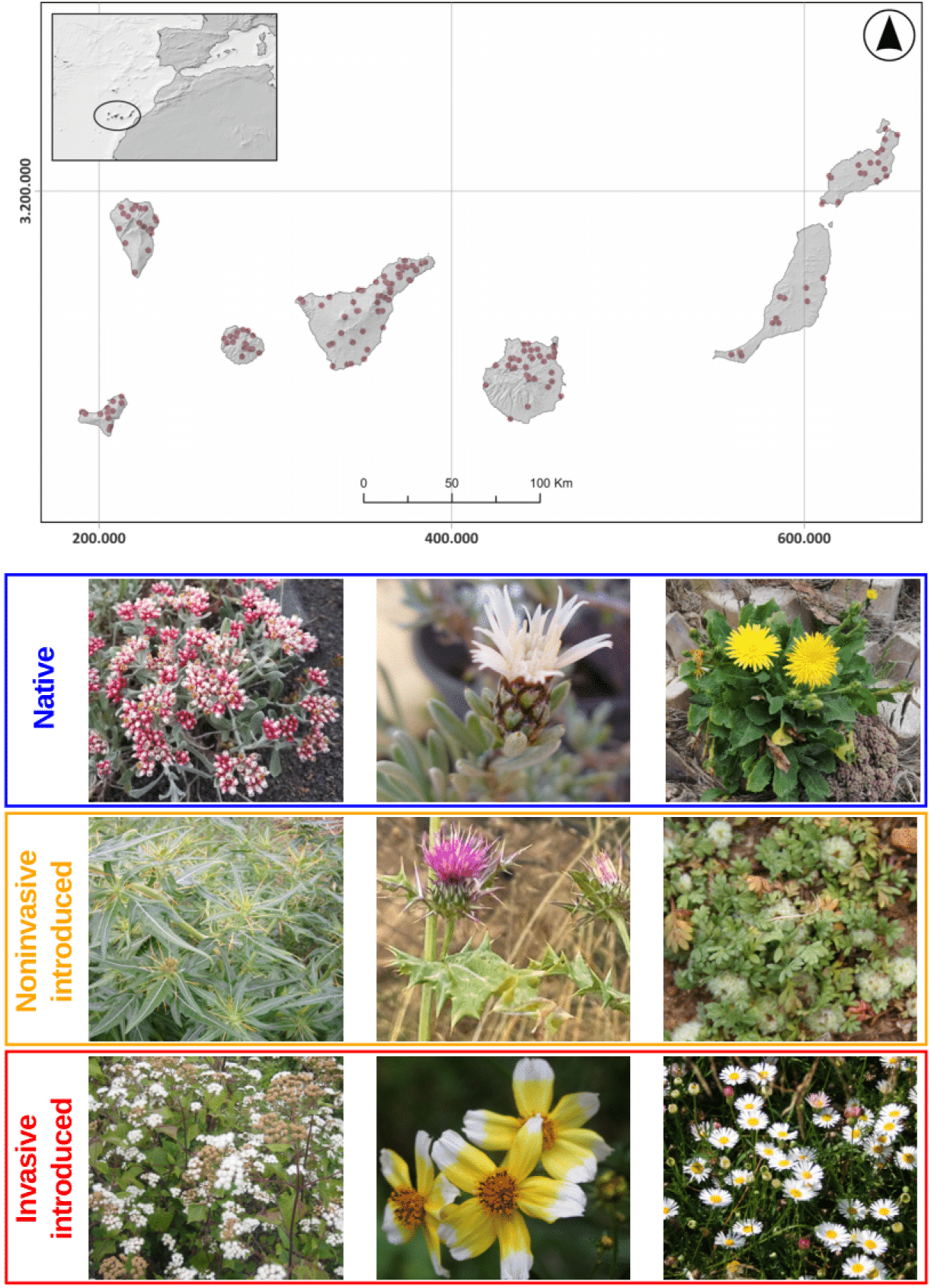
Map of the Canary Islands with all the sampling localities, and three representative species of each of the groups considered in this study: native, noninvasive introduced and invasive introduced species. From left to right and from top to bottom, the species within each group are: *Helichrysum monogynum*, *Atractylis preauxiana*, *Reichardia famarae* (natives); *Xanthium spinosum*, *Notobasis syriaca*, *Soliva stolonifera* (noninvasives); *Ageratina adenophora*, *Bidens aurea*, *Erigeron karvinskianus* (invasives).

When finding a species in the field was not possible, we visited the TFC and ORT herbaria to collect leaf samples from preserved specimens. These samples were used exclusively for DNA extraction, as leaf trait measurements require fresh material.

### DNA extraction and library preparation

We extracted total genomic DNA from silica-dried and herbarium leaf samples using commercial kits. For silica-dried fresh leaves, we used the Isolate II Plant DNA Kit (Bioline), the E.Z.N.A.^®^ Plant DNA DS Kit (Omega), and the Mag-Bind^®^ Plant DNA DS Kit (Omega).

The latter was used in combination with the ADN KingFisher Flex robotic system (ThermoFisher Scientific), applying half of the manufacturer’s recommended volume. For herbarium samples, we exclusively used the E.Z.N.A.^®^ Plant DNA DS Kit (Omega), as it demonstrated the best performance with this type of material. When the DNA extracts exhibited a greenish-brownish coloration, or had low DNA concentration (< 6–7 ng/µl), we cleaned-up and re-concentrated it using 2x Mag-Bind^®^ TotalPure NGS (Omega).

All DNA extracted from fresh samples was sheared before library preparation. We started enzymatically shearing the DNA by using NEBNext^®^ dsDNA Fragmentase^®^ (New England BioLabs), incubating samples at 37°C for 5–25 minutes. However, we soon realized that the optimal incubation time varied by species or, at best, by genus, making this method time- and resource-intensive. Consequently, we adopted mechanical shearing methods instead. For mechanical fragmentation, we processed 30 µl of genomic DNA per sample in a microTUBE-50 AFA Fiber Screw-Cap Case using a Covaris M220 Focused-ultrasonicator (Covaris, Woburn, Massachusetts, USA). To obtain fragment sizes around 300 bp, we applied the following settings: 65 s fragmentation time, 20% duty, 50 peak power and 200 cycles/burst. Fragment sizes from herbarium samples were assessed via 1% agarose gel electrophoresis. Those samples with already fragmented DNA or low DNA concentration (2–8 ng/µl) were directly used for library preparation, while the remaining samples were mechanically fragmented using the same ultrasonicator settings as fresh samples.

For library preparation, we used the NEBNext^®^ Ultra™ II DNA Library Prep Kit for Illumina^®^ (New England Biolabs), applying a half-volume protocol (Hale *et al*., 2020) with 200–300 ng of starting DNA whenever possible. We followed the manufacturer’s guidelines for selecting 300–400 bp fragments and uniquely labeled each sample using a combination of i7 and i5 primers from the NEBNext^®^ Multiplex Oligos for Illumina^®^ – Dual Index Primers Sets 1 and 2 (New England Biolabs), applying 10–15 cycles of PCR amplification. Dual-indexed libraries were pooled in groups of 10–28 samples based on DNA concentrations measured with a Qubit fluorometer (ThermoFisher Scientific, Inchinnan, UK). In no case were samples sheared with enzymatic and mechanical methods mixed in the same pool. Each pool underwent an independent hybridization reaction using a 8:1 mix of the commercial probes sets Angiosperms353 (Johnson et al., 2019; hereafter A353) and Compositae1061 (Mandel et al., 2014; hereafter COS), included in myBaits kits (ArborBiosciences). Probes were previously diluted in water at 1.5:1 ratio. The hybridization reaction was carried out at 65°C for 24 h. The enriched pools were amplified using KAPA HiFi HotStart Ready Mix (Roche) with 15 PCR cycles, followed by a 0.9x AMPure XP (Beckman Coulter) clean-up. The quality and average fragment size of the final products were assessed in a 2100 Bioanalyzer (Agilent). Enriched pools were multiplexed to contain between 16 to 188 libraries depending on the sequencing platform used. The enriched library pools were sequenced on either a MiSeq (v3, 2 x 300 bp paired-end reads) by the Instituto Tecnológico de Energías Renovables (ITER, Spain), or a HiSeqX (150 bp paired-end reads) and a Novaseq6000 (150 bp paired-end reads) by Macrogen (South Korea).

### Data processing

Raw sequences were quality filtered and adapters were removed using different software depending on the sequencing platform: Trimmomatic (Bolger et al., 2014) for reads from MiSeq and HiSeqX sequencing platforms, and fastp (Chen *et al*., 2018) for reads from Novaseq6000, following recommendations to account for differences in sequencing chemistry (M. Johnson pers. comm.). We set the parameters in the two softwares to obtain equivalent results. For Trimmomatic we used SLIDINGWINDOW:4:15, LEADING:3, TRAILING:3, and MINLEN:36, while in fastp we set the following equivalent parameters to similar values -r --cut_right_window_size 4 --cut_right_mean_quality 15, -5 --cut_front_window_size 1 --cut_front_mean_quality 3, -3 --cut_tail_window_size 1 --cut_tail_mean_quality 3, and -l 36.

The cleaned reads were used as input for HybPiper v. 2.0.2 (Johnson *et al*., 2016) to retrieve targeted genes. In brief, HybPiper maps reads to the targeted sequences, sorts them, and assembles contigs to extract exon and intron sequences separately. To construct the target sequence file, we concatenated reference sequences from the “mega353” (McLay *et al*., 2021) and the COS_sunf_lett_saff_all.fasta target files (https://github.com/Smithsonian/Compositae-COS-workflow/tree/master). To detect shared genes between the A353 and COS probe sets, we performed a BLAST search (Camacho et al., 2009) using the COS target file as database and the mega353 file as query. The genes from COS shared with A353 were removed from all the samples.

For those genes represented in at least 75% of the samples, we performed the sequence alignment of each gene separately. For the multiple sequence alignment we used MAFFT v7.511 with the “--auto” option (Katoh & Standley, 2013), and AMAS for the subsequent gene concatenation (Borowiec, 2016). Multiple sequence alignment and gene concatenation were performed independently with coding sequences (exon) and with exon+intron sequences (supercontigs), as recovered by HybPiper.

We selected three species as outgroups based on the backbone Asteraceae phylogeny (Mandel *et al*., 2019). These species were: *Gamocarpha macrocephala* S.Denham & Pozner (≡ *Nastanthus patagonicus* Speg.), belonging to the Calyceraceae family; and *Brunonia australis* Sm. ex R.Br. and *Scaevola tomentosa* Gaudich., both from the Goodeniaceae family. The raw data of these species were downloaded from the Sequence Read Archive (SRA) repository (see Supporting Information Table S3) and subjected to the same procedure as the raw reads generated for the present study, starting from HybPiper onwards. The only difference was the target file used in HybPiper, which depended on the specific probe set used to generate the outgroup data (mega353 or COS).

### Phylogenetic reconstruction and phylogenetic distances

To reconstruct the phylogenetic relationships, we used maximum-likelihood (ML) and multispecies coalescent (MSC) approaches using both exon and supercontig datasets. A ML phylogenetic tree was inferred using the concatenated multiple sequence alignment of all genes in IQTree v.2.3.0 (Santiago & Kim, 2009; Vitales *et al*., 2014; White *et al*., 2020) in independent runs for exon-only datasets and supercontig datasets. We used an unpartitioned scheme for which the appropriate substitution model was selected by ModelFinder (Kalyaanamoorthy *et al*., 2017). Branch support was estimated performing ultrafast bootstrap (Hoang *et al*., 2018) and SH-like approximate likelihood ratio test (Guindon *et al*., 2010), each of them with 1000 bootstrap replicates.

In addition, we used a MSC approach to infer a species tree summarizing individual gene trees. Gene trees were inferred with IQTree using the same parameters as the ML analysis, except that branch support was estimated only with ultrafast bootstrap. The resulting gene trees were used as input for ASTRAL-IV v.1.16.3.4 (Zhang & Mirarab, 2022) to infer the species tree under the MSC model. IQTree was used to estimate branch lengths on the ASTRAL tree topology obtained (‘-te’ option). This procedure was performed independently with both the exon-only and supercontig datasets.

We quantified the phylogenetic distances between introduced and native species using the independently generated phylogenetic ML and MSC trees based on the exon-only and supercontig datasets. For each described tree, we calculated two phylogenetic distance metrics: (i) Mean phylogenetic distance (MPD), and (ii) mean phylogenetic distance to the nearest native (MNND). The MPD is the average value of all mean phylogenetic distances between each introduced and all native species, while the MNND is the average value of the distances of each introduced species to its closest native species. These two phylogenetic distance metrics were calculated separately for noninvasive introduced species and invasive introduced species.

### Functional trait measurements and functional distance

Specific leaf area (SLA) and leaf dry matter content (LDMC) were measured following Cornelissen *et al*. (2003) and Pérez-Harguindeguy *et al*. (2013). In brief, we collected at least one leaf per individual from an average of three individuals per species in the field. In the lab, leaves were wrapped in soaked filter paper and stored overnight in zipped plastic bags at 4°C to maintain hydration. The next day, we recorded the water-saturated fresh mass of each leaf and scanned them for surface area measurements. Leaves were then oven-dried at 65°C until fully desiccated and weighed again to obtain dry mass. Leaf images were processed with cellSens Standard 3.1 Olympus software (Olympus Corporation, Tokyo, Japan) to get the leaf area. We calculated SLA by dividing leaf area by its dry mass (mm^2^ mg^-1^) and LDMC as leaf dry mass divided by its water-saturated fresh mass (mg g^-1^). Each species was functionally characterized by averaging all measurements for each leaf trait. In addition, the habit (woody *versus* herbaceous) for each species was recorded as a qualitative trait. To compare SLA and LDMC values across native, non-invasive, and invasive species, we conducted exploratory ANOVAs. Additionally, we used a Chi-squared test to assess differences in growth habit (i.e., woody *vs*. herbaceous) among these species groups.

We calculated multi-trait functional distances using the *gawdis* function from the “gawdis” R package (de Bello *et al*., 2021). This function calculates the distance by adjusting the weighting of each variable (or group of variables) to ensure an equal contribution to the overall distance metric. Following Götzenberger *et al*. (2020), we grouped all leaf traits into a single category since they were measured from the same organ. Two functional distances were calculated, following the same procedure described for the phylogenetic distances: (i) Mean functional distance (MFD) of each introduced to all native species, and (ii) mean functional distance to the functionally closest native species (MNFD). These functional distances were also calculated separately for noninvasive introduced species and invasive introduced species.

### Null models and standardized effect size

To evaluate whether introduced species are phylogenetically and functionally closer to or more distant from native species than expected by chance, we used null models. We first randomized the native/noninvasive/invasive species groupings 1000 times maintaining species richness in each group, and created 1000 null community matrices. For each matrix, we recalculated all phylogenetic and functional distances. We then compared the observed distance values to the distribution of null values to estimate the probability of obtaining the observed pattern by chance (*P*). If fewer than 5% of the null values were higher or lower than the observed value, we considered the result statistically significant (*P* < 0.05). We calculated the standardized effect size (SES) to quantify the deviation from the null expectation as:

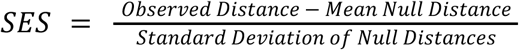

Positive SES values indicate that introduced species are more distantly related to natives than expected by chance, while negative SES values indicate that introduced species are functionally or phylogenetically closer to natives than expected by chance. To test the relative importance of the spatial scale in our analyses, we also calculated the functional and phylogenetic distances and performed the 1000 null models to calculate *P* and SES for each island independently. In addition, we compared all functional and phylogenetic distances calculated for invasive and noninvasive species at the two scales using t-tests.

## Results

### The Asteraceae family in the Canary Islands

The Biodiversity Data Bank of the Canary Islands (BIOTA) recorded 314 species of Asteraceae in February 2024, including 227 native species (138 endemics) and 87 introduced species (9 invasive). After reviewing this checklist, we excluded 21 species due to misidentifications, uncertain records, or lack of naturalized occurrence confirmations. In addition, seven species recently added in 2024 to the checklist were not included in our study, and 20 species could not be found in the field or herbaria due to their rarity or difficult identification. Consequently, our study focused on 266 species recorded in BIOTA, comprising 213 native species and 62 introduced species, plus nine not recorded in BIOTA (newly described or previously undocumented during the present study; Supporting Information Table S1, 275 species in total). Most species were sampled in the field, allowing for both DNA isolation and trait measurements, while 26 species were only sampled from herbaria (ORT and TFC) for DNA isolation.

### Targeted gene sequences

Out of the 275 samples collected for DNA isolation and library preparation, seven samples failed during the wet-lab procedure. From the remaining 268 samples, we obtained, on average, 5.7 millions of reads per sample (min. 459,250 reads), with a target enrichment efficiency (i.e., number of reads mapped to the target loci / total number of reads) of 61.5% (12.7–85.7%) . The number of exons recovered ranged from 414 to 1391 genes per sample (mean = 1250.88 genes/sample), while supercontig recovery ranged from 603 to 1396 genes per sample (mean = 1265.80 genes/sample) (Supporting Information Table S2). Of the targeted 1414 loci, 1412 were successfully recovered across all samples. Specifically, the A353 probe set recovered 352 genes, and the COS probe set recovered 1060 genes. After excluding 39 overlapping genes and those with coverage in fewer than 75% of samples, 1187 genes were retained (329 from A353 and 858 from COS probe sets, respectively).

### Phylogenetic inference and incongruence

The phylogenetic relationships within the three Asteraceae subfamilies in the Canary Islands were well resolved and consistent across both phylogenetic methods and datasets (Fig. 2). In general, we obtained consistent topologies and good node support values, with some exceptions. Within the subfamily Asteroideae, the sister tribes Gnaphalieae (*Filago*, *Gamochaeta*, *Helichrysum*, *Ifloga*, *Laphangium*, *Phagnalon*) and Astereae (*Erigeron*, *Bellis*, *Symphyotrichum*), are sister to the tribes Senecioneae (*Pericallis*, *Delairea*, *Kleinia*, *Bethencourtia*, *Senecio*, *Roldana*) and Anthemideae (*Argyranthemum*, *Otoglyphis* [≡ *Aaronshonia*], *Achillea*, *Anacyclus*, *Anthemis*, *Artemisia*, *Cladanthus*, *Coleostephus*, *Cotula*, *Glebionis*, *Gonospermum*, *Matricaria*, *Soliva*) in the ML tree built using exon sequences; while the tribe Senecioneae is retrieved as basal in this four-tribe group in the MSC tree topology (exon-only and supercontig) and the ML tree with supercontig sequences. In the subfamily Cichorioideae, we observed similar discrepancies in the clades grouping certain genera. Specifically, *Andryala*, *Cichorium*, and *Tolpis* form a clade that is more basal in the ML trees (exon-only and supercontig) and the MSC tree built with the supercontig dataset. Conversely, the clade containing *Crepis*, *Hedypnois*, *Helminthotheca*, *Hypochaeris*, *Picris*, *Rhagadiolus*, *Scorzoneroides*, and *Thrincia* occupies a more basal position in the MSC species tree with the exon-only dataset. Additional inconsistencies were observed in the phylogenetic positions of some recently radiated genera such as *Sonchus*, *Cheirolophus* and *Argyranthemum*; as well as in the position of the genus *Onopordum* within the subfamily Carduoideae (Supporting Information Figs. S1–S4).

**Figure 2.**
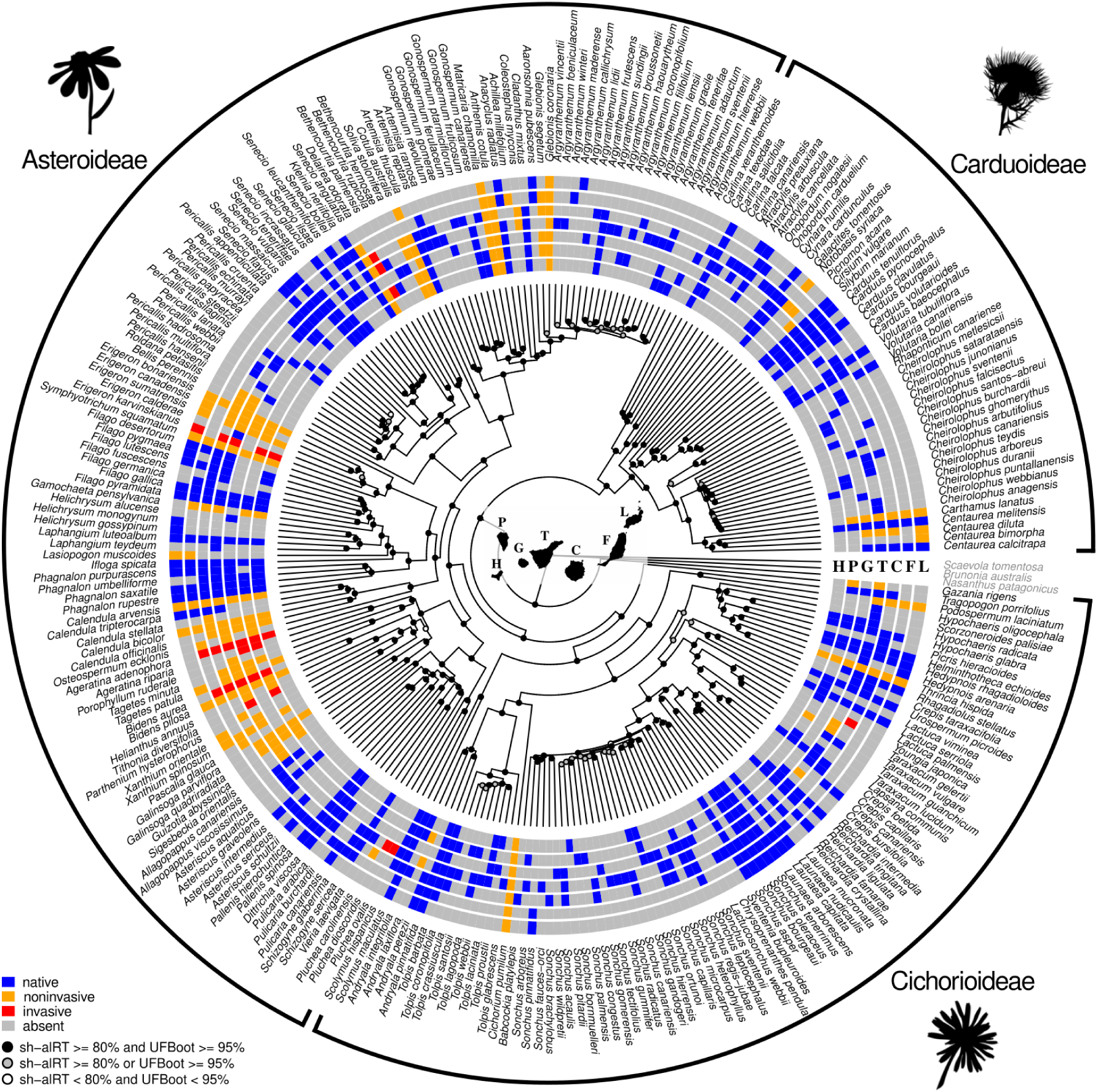
Maximum likelihood phylogenetic tree estimated with IQTree from exon-only sequence data. Node support is indicated with circles. Solid black circles indicate full support in both sh-alRT and UFBoot tests, grey circles indicate support in only one of the tests, and white circles indicate statistically not supported clades. Species distribution is shown in seven concentric colored circles: blue indicates the presence of native species; orange, noninvasive introduced species; red, invasive introduced species; and grey indicates the absence of the species on an island. Island abbreviations: H, El Hierro; P, La Palma; G, La Gomera; T, Tenerife; C, Gran Canaria; F, Fuerteventura; L, Lanzarote. The three main subfamilies present in the archipelago are also indicated. Outgroup species are shown in grey. Accepted names of the synonyms used in the figure: *Gamocarpha macrocephala* (≡ *Nastanthus patagonicus*), *Otoglyphis pubescens* (≡ *Aaronshonia pubescens*). Plant silhouettes are from PhyloPic (www.phylopic.org).

### Functional traits

Leaf functional traits were measured for 228 species, representing 85% of the species included in the phylogeny (see Supporting Information Fig. S5 and Table S4). Of these, 181 were native to the Canary Islands, 40 species were noninvasive introduced and seven were invasive introduced species. Exploratory ANOVAs comparing SLA and LDMC values between natives, noninvasive and invasive species revealed no significant differences between groups for either trait (SLA: *F* = 0.876, *P* = 0.418; LDMC: *F* = 0.794, *P* = 0.453). Regarding the woody/herbaceous habit of the species, we found that most invasive and native species were woody, while noninvasive introduced species tended to be predominantly herbaceous. Indeed, both variables, the woody/herbaceous habit and the origin of the species, were significantly associated (*χ*^2^ = 32.463, *P* = 0.001).

### Testing Darwin’s naturalization conundrum at different spatial scales

At the archipelago scale, both noninvasive and invasive introduced species were found to be phylogenetically distant from the entire native community and from their closest native relative, compared to the null expectation (SES > 0). This pattern was consistently significant for noninvasive and invasive introduced species in relation to their closest native relative (*P* < 0.05) (Fig. 3A, B). The distant relationship between noninvasive species and the entire native community was significant. However, the distance between invasive species and the entire native community was not statistically supported when using the exon-only ML and MSC trees (Fig. 3A, Supporting Information Table S5).

**Figure 3.**
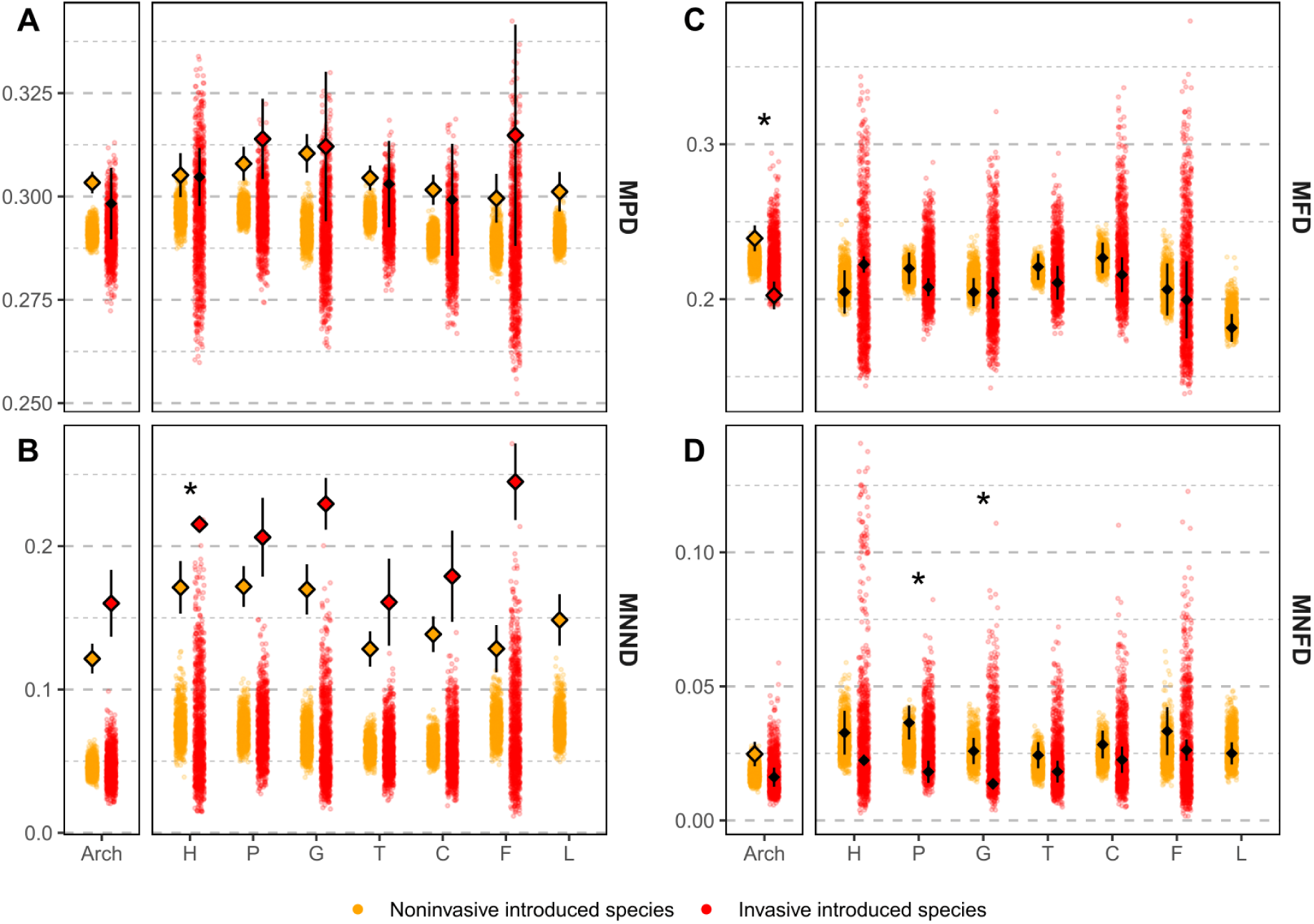
Phylogenetic (A, B) and functional (C, D) distance metrics between noninvasive (orange) and invasive (red) introduced species. Upper panels show the mean phylogenetic/functional distance of each introduced species to all the natives (MPD – A, and MFD – C), while lower panels show the mean phylogenetic/functional distance of each introduced species to its closest native (MNND – B, and MNFD – D). The point clouds indicate all null values obtained by randomly reshuffling the community matrices 1000 times. When the observed value was significantly higher or lower than expected by chance (SES > 0 or SES < 0), it is indicated with a colored diamond, while it is shown as a solid black diamond when no statistical support was obtained. Standard error bars extend from the diamond and indicate the standard error of all phylogenetic distance measures of each introduced species to either the whole native community or the closest native species. When difference between noninvasive and invasive species were statistically supported, it is indicated with asterisks (*: *P* < 0.05). Arch: archipelago, H: El Hierro, P: La Palma, G: La Gomera, T: Tenerife, C: Gran Canaria, F: Fuerteventura, L: Lanzarote.

At the island level, we obtained similar results. Introduced species were phylogenetically distant from both the whole native community and their closest native relatives across all islands, when compared to the null expectation (SES > 0). This result was statistically supported for noninvasive and invasive introduced species to their closest native relative. Invasive species showed a statistically significant distant relationship with the whole native community on five of the six islands with more than one invasive species when using the supercontig ML and MSC trees (El Hierro, La Palma, La Gomera, Tenerife, Fuerteventura; Supporting Information Table S5), and on only three islands when using the exon-only trees (La Palma, La Gomera, Fuerteventura; Fig. 3A, Supporting Information Table S5). At both scales, when considering the mean distance of introduced species to their closest relative, we obtained higher values for invasive species than for noninvasive introduced species (Fig. 3B). However, in most cases this difference was not statistically supported.

At the archipelago scale, we observed contrasting results for noninvasive and invasive introduced species in terms of multi-trait functional distances. Noninvasive introduced species were functionally more distinct from the entire native community and from their closest native relative than expected by chance (SES > 0). Surprisingly, invasive introduced species were functionally similar to the native community and to their closest native relative (SES < 0). This trend was statistically supported for all comparisons, except for the distance between invasive species and their functionally closest native relative (MNND_SES_ = -0.218, *P* = 0.475) (Fig. 3C, D). At the island scale, a similar pattern was observed, with some exceptions, although none of the results reached statistical significance at this level (Supporting Information Table S6).

When comparing functional distances to natives (MFD and MNFD) between noninvasive and invasive introduced species, we detected a general tendency of invasive species being functionally more similar to both the whole native community and their functionally closest native species than noninvasive species (Fig. 3C, D). This difference was statistically significant at the archipelago level for MFD (*t* = 2.999, *d.f.* = 18.565, *P* = 0.008), and for two of the six islands with more than one invasive species for MNFD (La Palma: *t* = 2.423, *d.f.* = 27.104, *P* = 0.022; La Gomera: *t* = 2.332, *d.f.* = 20.752, *P* = 0.030).

## Discussion

Our results show that noninvasive and invasive Asteraceae species introduced to the Canary Islands are phylogenetically distant from native species, with invasive species exhibiting slightly greater phylogenetic divergence from their closest native relatives than noninvasive ones. This pattern aligns with Darwin’s naturalization hypothesis, which suggests that biotic filtering favors the establishment of more distantly related species to minimize competitive interactions. Accordingly, we found that noninvasive species also show functional divergence from native counterparts. However, invasive introduced species display a contrasting trend, being functionally more similar to native species than expected by chance and, therefore, providing partial support to the pre-adaptation hypothesis. These findings, consistent across multiple phylogenetic reconstructions and spatial scales, underscore the complexity of the invasion dynamics (Theoharides & Dukes, 2007; Gioria *et al*., 2023).

Darwin’s naturalization conundrum has been extensively tested, often yielding seemingly contradictory results (reviewed in Thuiller *et al*., 2010; Cadotte *et al*., 2018). However, these discrepancies can be attributed to the influence of multiple factors and the intricacy of ecological processes at play. Recent studies have contributed to reconciling the two competing hypotheses by incorporating broader ecological and evolutionary perspectives (Schaefer *et al*., 2011; Li *et al*., 2015; Omer *et al*., 2022; Fan *et al*., 2023; Le *et al*., 2024). In the present study, we found evidence for both of Darwin’s hypotheses, revealing a decoupling between phylogenetic and functional distances to natives when comparing invasive and noninvasive plant species within the Asteraceae family.

### Patterns and drivers of phylogenetic and functional dissimilarity

Oceanic island floras are often phylogenetically clustered and disharmonic compared to their continental counterparts (Weigelt *et al*., 2015; Taylor *et al*., 2019; König *et al*., 2021). Species colonizing oceanic islands must first overcome the dispersal barrier imposed by the surrounding ocean. Once established, some lineages undergo extensive diversification contributing to filling the numerous available niches in these newly emerged, initially lifeless territories (Patiño *et al*., 2014; Whittaker *et al*., 2023). As a result, the native island floras often exhibit large phylogenetic gaps, which may favour the naturalization of distantly related introduced species (Bach *et al*., 2022; Le *et al*., 2024). This pattern is consistent with the positive MPD_SES_ values observed in our study for both noninvasive and invasive introduced species, as MPD is a distance metric sensitive to deep phylogenetic branching and reflects rather old phylogenetic relationships (Mazel *et al*., 2016). Moreover, introduced species were also distantly related to their closest native relatives (MNND_SES_ > 0), suggesting a consistent pattern across both deep and recent evolutionary timescales. This result is typically interpreted as an outcome of competitive interactions that hinder the establishment of closely related species. Nevertheless, competition may result in different phylogenetic outcomes and its role should be further assessed using functional traits (Cahill *et al*., 2008; Thuiller *et al*., 2010; Gioria *et al*., 2023; Hähn *et al*., 2025).

Leaf functional traits did not differ significantly between introduced and native species, a somewhat unexpected result. For instance, the specific leaf area, which is positively related with the growth rate, has been associated with successful invasive species (Pyšek & Richardson, 2007). However, in this study, the only significant differences among invasive, noninvasive and native species were found in their herbaceous *versus* woody habit. Noninvasive introduced species were predominantly herbaceous plants, whereas most invasive and native species had a woody habit. Although herbaceous plants typically have short life cycles and efficient dispersal capabilities, which can be advantageous for invasiveness (Cadotte & Lovett-Doust, 2001; Moravcová *et al*., 2015), some studies suggest that the dominant life form among nonnative floras are stage-specific, with annuals more common in early invasion stages and successful invaders more often being long-lived species (Pyšek & Richardson, 2007). This is in agreement with the findings from the Azorean flora, where herbaceous plants had lower probability of becoming invasive compared to other life forms (Schaefer *et al*., 2011). Ultimately, invasion success is also influenced by the characteristics of the recipient community and how different the introduced species are from it (Catford *et al*., 2019). Divergence between native and introduced species is at the heart of Darwin’s hypotheses, and its effect on invasion success sometimes exceeds the effect of individual trait values alone (Carboni *et al*., 2016; Catford *et al*., 2019).

The estimated multi-trait distance yielded contrasting patterns depending on the invasion stages. Noninvasive introduced species were functionally distant from native species, whereas invasive species were functionally similar. Functional similarity between invasive and native species has been reported in several studies. For example, Schaefer *et al*. (2011) found a correlation between invasion probability and similarity in life form between invasive and native species across the Azores. Similarly, invasive species in the San Juan Islands exhibited functional resemblance to natives, although the strength of this pattern varied depending on the trait assessed (Marx *et al*., 2016). Le *et al*. (2024) observed that in flooded areas, invasive species were phylogenetically distant from natives but functionally similar. Likewise, Park *et al*. (2024) found that among California’s angiosperms, invasive species exhibited high functional similarity in flowering phenology despite being phylogenetically distant from natives. This decoupling between phylogenetic and functional distances to native species has been attributed to a combination of competition and environmental filtering. While invasive species that are phylogenetically distant from natives can exploit unoccupied evolutionary space avoiding competition with close relatives, strong environmental filtering selects for species with functional traits similar to those of native species (Le *et al*., 2024). Another possible explanation is the ‘enemy release’ hypothesis, which predicts that distantly related introduced species will succeed due to the absence of natural enemies in the recipient community, regardless of functional similarity to native species (Schaefer *et al*., 2011; Bezeng *et al*., 2015).

Interpreting the mechanisms underlying phylogenetic-functional relationships is further complicated by studies reporting the opposite pattern of phylogenetic-functional decoupling, where introduced species are phylogenetically close but functionally distant from native species (Ordonez, 2014; Marx *et al*., 2016). Recent evidence highlights the importance of the phylogenetic and functional decoupling in community assemblages, with evidence for the two opposing patterns, and conclude that phylogenetically distant species often share similar functional traits, while closely related species may be functionally different (Večeřa *et al*., 2023; Hähn *et al*., 2025). While our results provide significant insights into the functional distance between introduced and native species, we acknowledge that key traits relevant to invasion success, such as reproductive strategies, plant height, nutrient content, and plant phenology were not included (Pyšek & Richardson, 2007; Schaefer *et al*., 2011; Ordonez, 2014; Bezeng *et al*., 2015; Carboni *et al*., 2016; Marx *et al*., 2016; Park *et al*., 2024). Furthermore, differences in less easily quantifiable but relevant traits in terms of competition, such as herbivory resistance, may remain hidden when using conventional functional traits (Hähn *et al*., 2025). Given that individual traits can yield contrasting results when analysed separately (Schaefer *et al*., 2011; Marx *et al*., 2016), incorporating a broader and more diverse set of traits would provide a more comprehensive and robust multi-trait measure that reliably reflects the functional mechanisms shaping invasion dynamics.

### The role of the invasion, spatial and evolutionary scales

The scale of invasion plays a key role in reconciling Darwin conundrum’s two hypotheses, as their support shifts depending on the stage considered (Cadotte *et al*., 2018; Omer *et al*., 2022). Our findings show that invasive species tend to be even more distant from their nearest native relatives than noninvasive species, yet they exhibit greater functional similarity to native species. Similarly, invasive species were more distantly related to natives than noninvasive species in continental regions across South Africa (Bezeng *et al*., 2015; Omer *et al*., 2022), California (Strauss *et al*., 2006), and oceanic archipelagos like the Azores (Schaefer *et al*., 2011). However, this result contrasts with the temporal study of Li *et al*. (2015), which tracked 480 plots during 40 years. They found that invasion success was negatively correlated with phylogenetic distance, i.e., introduced species with close native relatives were more likely to establish. This study also revealed a high extinction probability for these closely related natives, ultimately resulting in communities dominated by invasive species coexisting with distantly related natives. Although this may occur at a small scale (i.e., plot level), it is less likely to hold at larger scales, where native species have greater chances of persisting (Carboni *et al*., 2016). In the Canary Islands, for example, recorded extinctions of Asteraceae species have been limited to four genera well represented in the native flora of the archipelago (Orihuela-Rivero *et al*., 2025). Notably, only one of them includes a noninvasive introduced species within the Canaries. The absence of closely related invasive counterparts among these extinction records, coupled with the relatively broader spatial scale of our study, supports the idea that our findings align with the temporal dynamic of the invasion process.

The spatial scale of the study is indeed another widely recognised factor influencing the outcomes of Darwin’s naturalization conundrum (Thuiller *et al*., 2010; Carboni *et al*., 2016; Marx *et al*., 2016; Cadotte *et al*., 2018). At smaller scales, competition and abiotic pressures, such as local disturbances or specific microclimates, tend to be stronger (Thuiller *et al*., 2010; Marx *et al*., 2016; Le *et al*., 2024), while larger spatial scales encompass greater environmental heterogeneity and more complex biotic and abiotic interactions (Thuiller *et al*., 2010; Marx *et al*., 2016). In our study, we observed no substantial differences between the archipelago-wide and island-level scales. While some standardized effect sizes of functional metrics showed opposite trends at the island scale compared to the entire archipelago, these trends lacked statistical support (Supporting Information Table S6). Bach *et al*. (2022) found that native phylogenetic clustering positively correlated with introduced species richness, particularly on smaller islands, consistent with Darwin’s naturalization hypothesis. In contrast, the pre-adaptation hypothesis was more likely to hold on larger islands (Bach *et al*., 2022). We found no such effect, possibly due to our focus on a single family and only seven islands, where even smaller islands contain highly heterogeneous habitats, which may be even more determinant than island size *per se* (Thuiller *et al*., 2010; Cadotte *et al*., 2018; Essl *et al*., 2019).

A limitation of our study is the lack of species coexistence data, making it unsuitable for resolving invasion dynamics within communities (Thuiller *et al*., 2010; Ng *et al*., 2019). However, broad-scale studies remain valuable, as species distributions reflect both local and global-scale processes (Park *et al*., 2020; Fan *et al*., 2023). Furthermore, our phylogenetic approach also ensured accurate reconstruction of evolutionary relationships within the family, often difficult in whole-community studies (Park & Potter, 2013; Omer *et al*., 2022; Fan *et al*., 2023). In fact, phylogenetic scale may also explain discrepancies between studies (Thuiller *et al*., 2010; Ng *et al*., 2019). By focussing on a single family, we assessed invasion dynamics among close relatives, and what appears phylogenetically distant at this scale may be considered closely related in broader community-wide analyses. A logical next step would be to analyze the entire Canarian flora to assess the consistency of our findings at a broader evolutionary scale.

### Concluding remarks

In conclusion, our study is the first to test Darwin’s naturalization conundrum in the most speciose angiosperm family of the Canary Islands, a biodiversity hotspot facing urgent global change challenges and threats (Morente-López *et al*., 2023; Patiño *et al*., 2023; Orihuela-Rivero *et al*., 2025). By constructing a highly resolved phylogeny encompassing most species in the archipelago, we highlight that introduced Asteraceae species distantly related to natives are more likely to become invasive. We emphasize the need to assess both phylogenetic and functional relationships, as phylogenetic evolutionary relatedness is not always a good proxy for ecological similarity (Ordonez, 2014; Marx *et al*., 2016; Cadotte *et al*., 2018; Bach *et al*., 2022). Future research should focus on species coexistence at broad taxonomic scales to further unravel invasion dynamics in oceanic archipelagos.

## Supporting information

SupportingInformation_Figures

SupportingInformation_Tables

## Acknowledgements

We would like to thank all the people who contributed in some way to this study. Matt Johnson and Lisa Pokorny for their guidance through the target enrichment protocol. Stephan Scholz, Aurelio Acevedo, Sébastien Mirolo, Dagmar Hanz, Marco Díaz-Bertrana Sánchez, Alicia Martín Alonso, Carlos Cáceres Marrero and Rubén C. García Medina for their contribution to field sampling. Cristina González Montelongo for her assistance at TFC herbarium. Antonio Íñigo Campos, Marina Ventayol and Antonio Pérez Delgado for their assistance and/or advice in the molecular wet lab. Abraham Araña Padilla, Carmen Balibrea Escobar and all students that have collaborated in the iEcoEvoLab group during the lifespan of the project (Annika Jurda, Mario Martín Almeida, Simon Mazaud, Carlos de la Rosa Báez, María Ruiz Navarro and Valentina León Pérez). This research was supported by the Spanish Ministry of Science and Innovation (MICINN) projects ASTERALIEN (PID2019-110538GA-I00) and DecodAdapt (PID2023-147122NB-I00), and the Fundación BBVA project INVASION (PR19_ECO_0046). JP also received funding from the MICINN through the Ramón y Cajal Program (RYC-2016-20506). LSJ-G was co-funded by the Academia Canaria de Investigación, innovación y Sociedad de la Información de la Consejería de Universidades, Ciencia e Innovación y Cultura and by European Social Fund Plus (ESF+) (FPI 2021 Fellowship TESIS2021010101). RO-R was supported by the University Teacher Training (FPU) program (FPU21/02585) funded by the Ministry of Universities. JM-L was funded by a Juan de la Cierva-Formación Fellowship (MICIIN; FJC2020-046353-I). We thank Teide High-Performance Computing facility (TeideHPC) of the Instituto Tecnológico y de Energías Renovables (ITER S.A.), for providing access to computer resources. We also thank all the public administrations and their personnel for the help and providing sampling permits: Gobierno de Canarias (2020/3987, 2022/602), Cabildo de El Hierro (4862/2021), Cabildo de La Palma (CA/00000110/0001/000038829), Cabildo de Tenerife (AFF 56/21), Cabildo de Gran Canaria (FLA 61-2022), Cabildo de Fuerteventura (2021/00018153H), Cabildo de Lanzarote (2021-1434), PN Caldera de Taburiente (APM/MFM), PN Garajonay (14.077), PN El Teide (MDV/amp), PN Timanfaya (RES-AUT I08/2021), PN Isla de La Graciosa (INV-02-20201).

## Author contributions

JP designed the research with inputs from YA and LSJ-G. JP, YA, LSJ-G, JAR-B, JM-L, GS-P, MAP-M, RO-R, AN-C, MS-P collected leaf samples. YA, LSJ-G, JM-L, GS-P, RO-R measured functional traits. YA performed the molecular wet lab, with advice of RG-M, and with LSJ-G and GS-P contributing with DNA extractions. YA analysed and interpreted the data with the help of JM-L and JP. YA and JP wrote the first version of the manuscript, which was improved by contributions from all authors.

## Data availability

Scripts are stored and publicly available in figshare repository (DOI: 10.6084/m9.figshare.28762889). Raw sequences will be available on NCBI SRA upon acceptance.

