## SupportingInformation_Figures for "Taming Darwin’s conundrum: On the role of phylogenetic and functional differentiation in plant naturalization on oceanic islands"

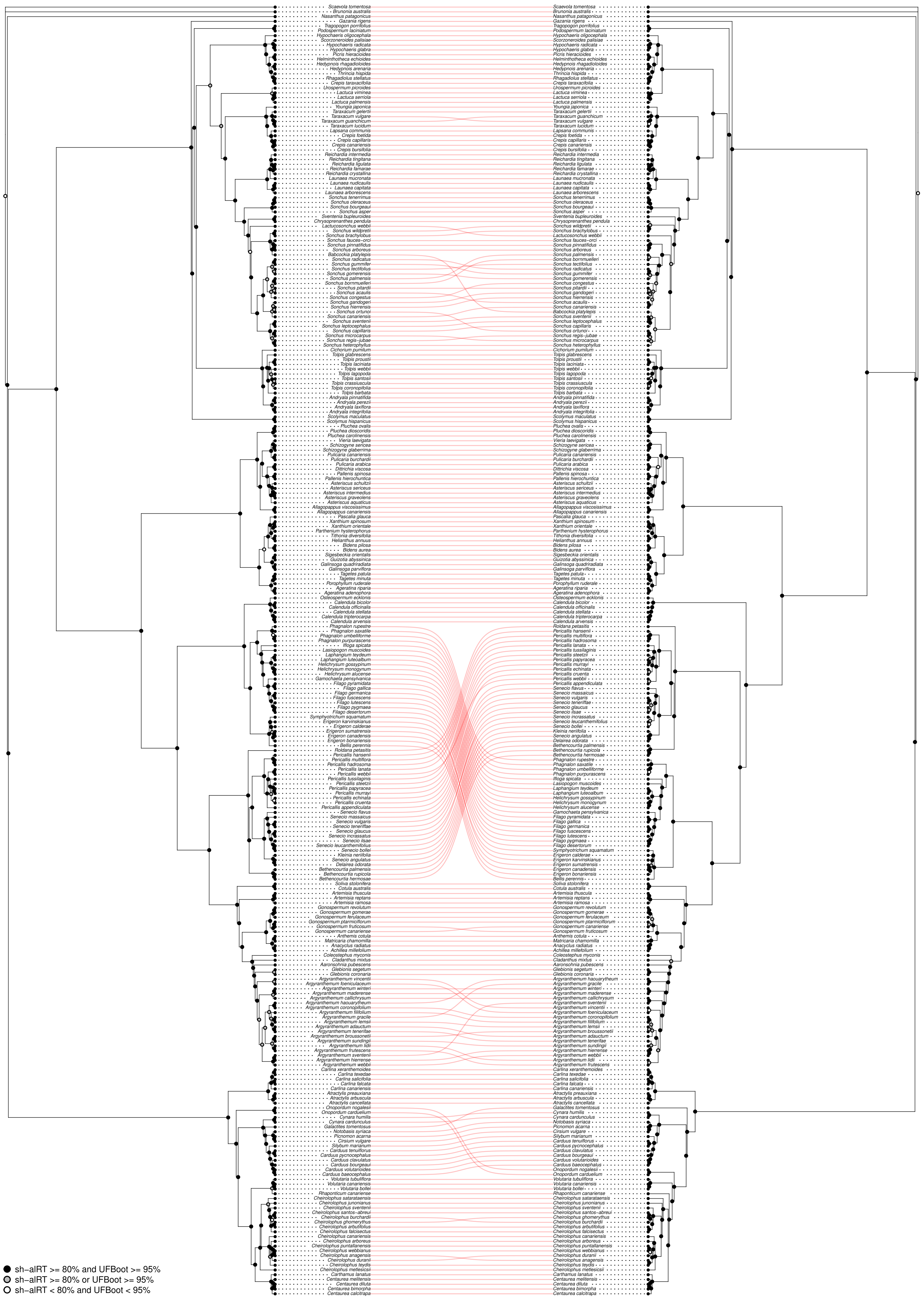


**Figure S1.** Comparison of maximum likelihood trees built with exon-only (left) and supercontig (right) sequences. Node support is indicated with circles: solid black circles indicate full support in both sh-alRT and UFBoot tests, grey circles indicates support in only one of the tests, and white circles indicate statistically not supported clades. Accepted names of the synonyms used in the figure: *Gamocarpha macrocephala* ( ≡ *Nastanthus patagonicus*), *Otoglyphis pubescens* ( ≡ *Aaronshonia pubescens*).

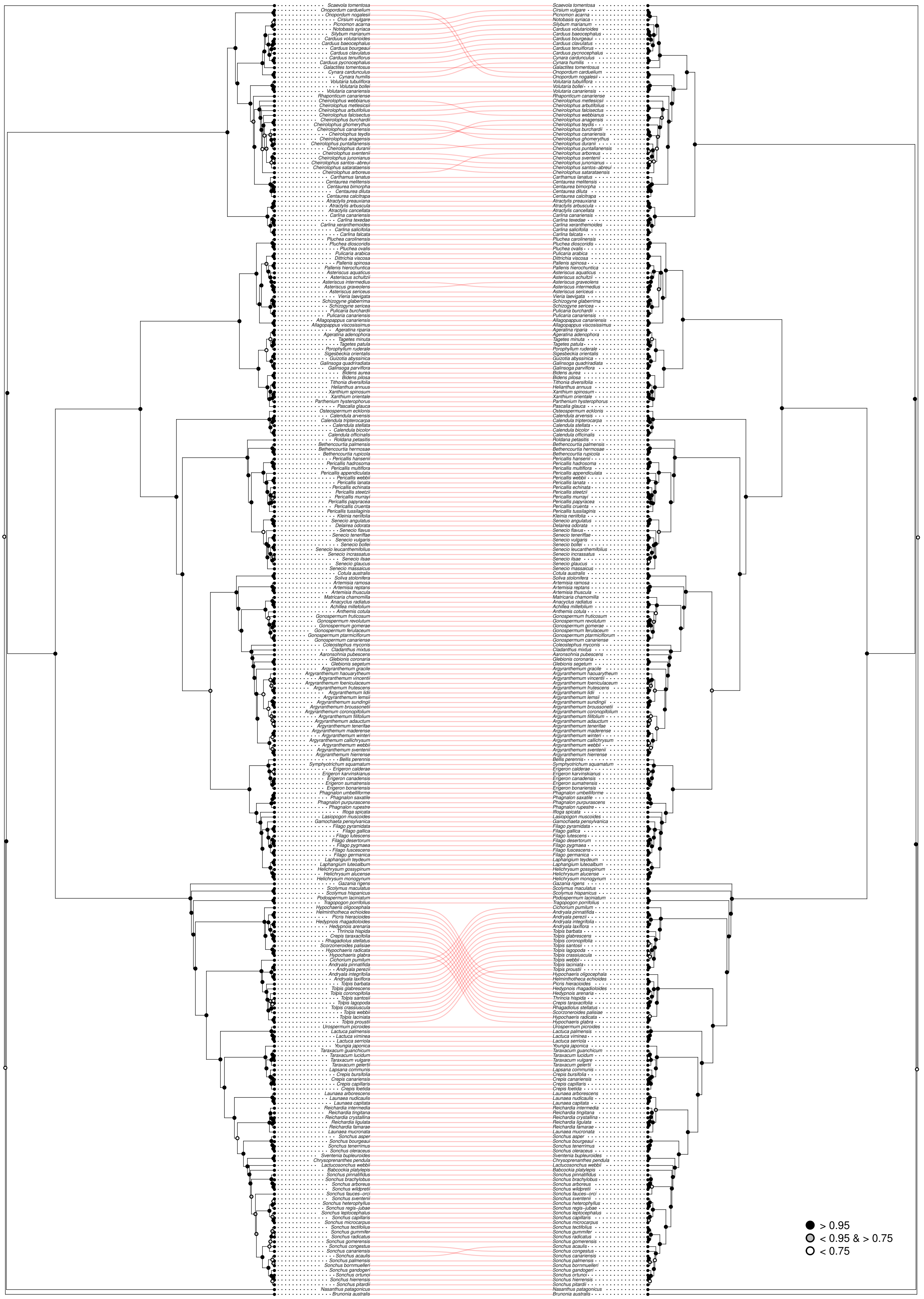


**Figure S2.** Comparison of multispecies coalescent trees (ASTRAL) built with exon-only (left) and supercontig (right) sequences. Node support is indicated with circles based on local posterior probabilities (see legend). Accepted names of the synonyms used in the figure: *Gamocarpha macrocephala* (≡ *Nastanthus patagonicus*), *Otoglyphis pubescens* (≡ *Aaronshonia pubescens*).

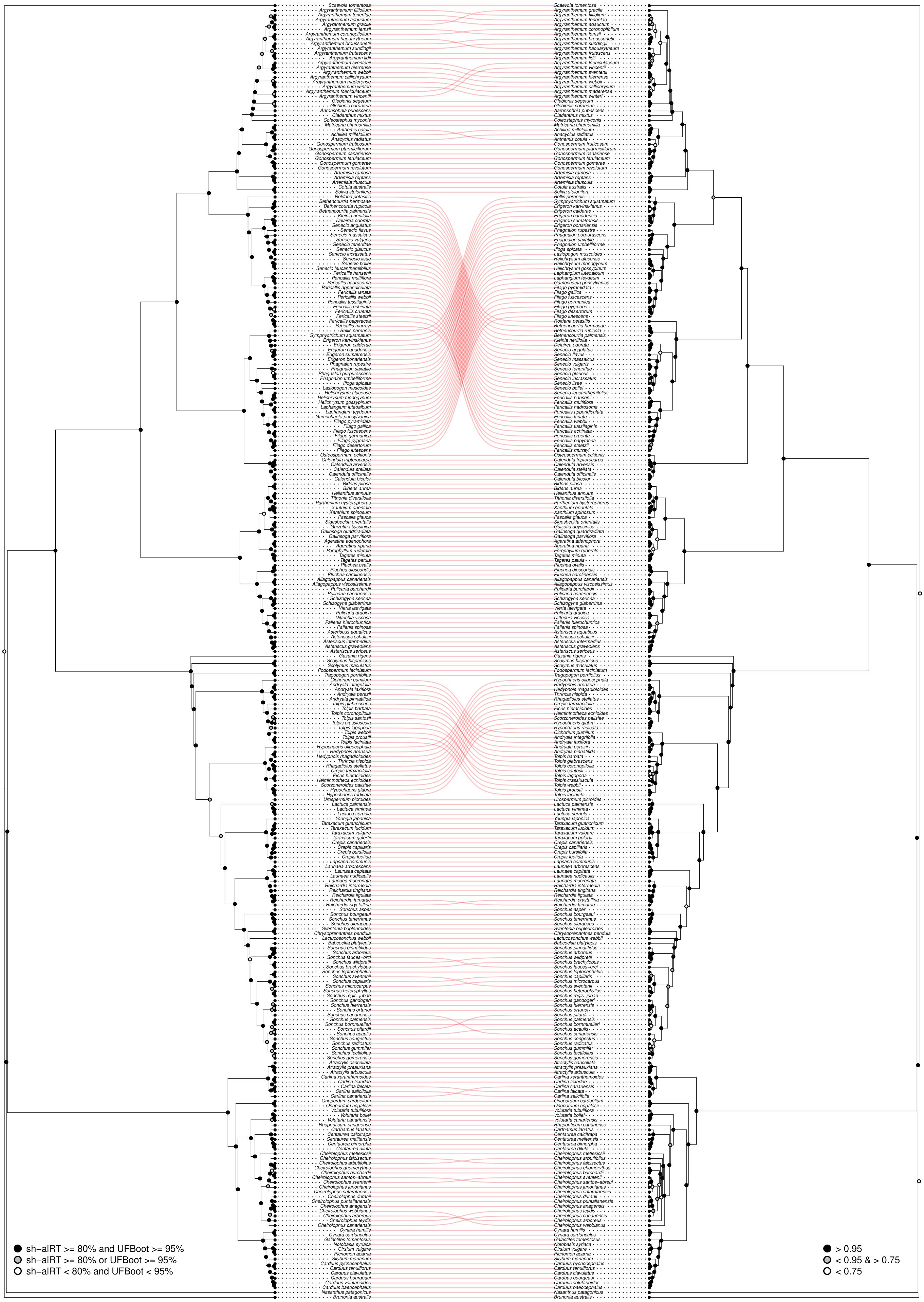


**Figure S3.** Comparison of maximum likelihood (left) and multispecies coalescent trees (ASTRAL, right) built with exon-only sequences. Node support is indicated with circles based on sh-alRT and UFBoot tests for the ML tree and on local posterior probabilities for the MSC tree (see legend). Accepted names of the synonyms used in the figure: *Gamocarpha macrocephala* ( ≡ *Nastanthus patagonicus*), *Otoglyphis pubescens* ( ≡ *Aaronshonia pubescens*).

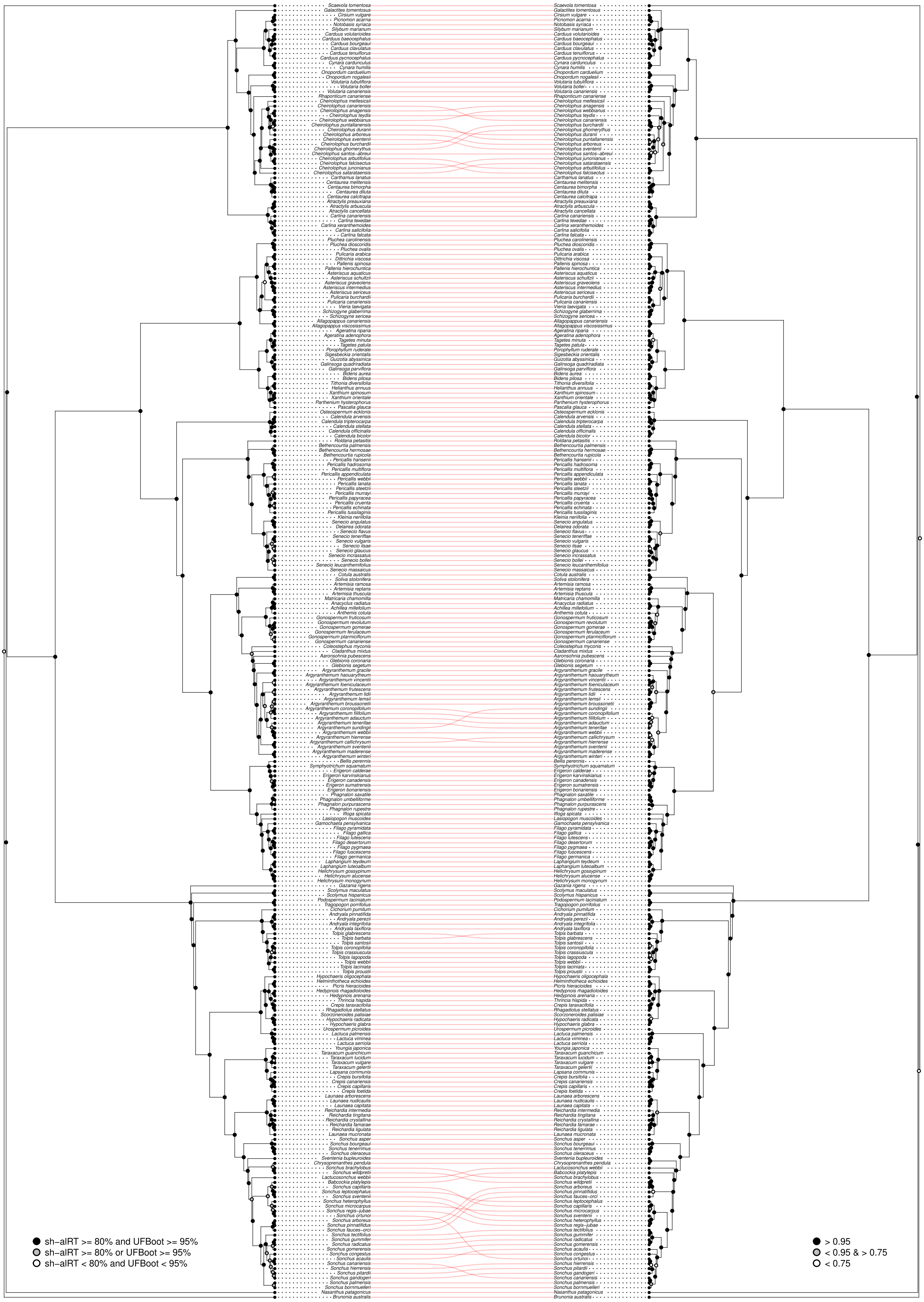


**Figure S4.** Comparison of maximum likelihood (left) and multispecies coalescent trees (ASTRAL, right) built with supercontig sequences. Node support is indicated with circles based on sh-alRT and UFBoot tests for the ML tree and on local posterior probabilities for the MSC tree (see legend). Accepted names of the synonyms used in the figure: *Gamocarpha macrocephala* ( ≡ *Nastanthus patagonicus*), *Otoglyphis pubescens* ( ≡ *Aaronshonia pubescens*).

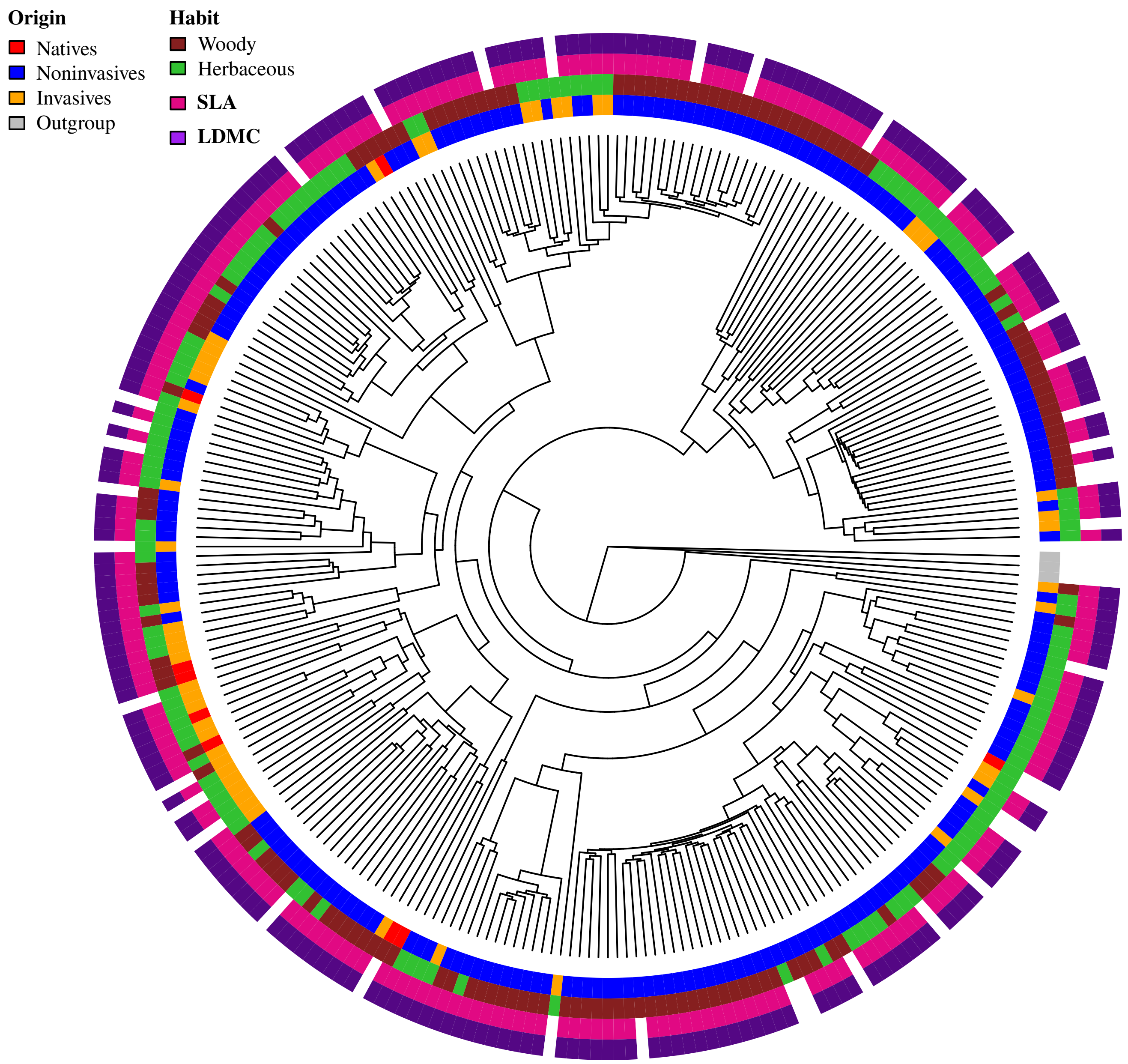


**Figure S5.** Maximum likelihood phylogenetic tree built with exon-only sequences. The four outer concentric circles indicate, from inside to outside: (1) the native (blue), noninvasive introduced (orange) or invasive introduced (red) origin of the species, the outgroup is indicated in grey; (2) the woody (brown) or herbaceous (green) habit of the species; (3) the species with measured SLA (pink); and (4) the species with measured LDMC (purple).
